# Whole-genome sequencing reveals inter-household networks of gut-colonising ESBL-producing *Escherichia coli* in two rural Malawian districts

**DOI:** 10.64898/2026.02.11.705350

**Authors:** Angus M. O’Ferrall, David Lally, Peter Makaula, Gladys Namacha, Joseph M. Lewis, Patrick Musicha, Richard N. Goodman, Ellie Allman, Sabrina Moyo, Claire S. Waddington, Sekeleghe A. Kayuni, Nicholas A. Feasey, Janelisa Musaya, J. Russell Stothard, Adam P. Roberts

**Author notes:** Corresponding author –.

## Abstract

Infection with extended-spectrum beta-lactamase-producing *Escherichia coli* (ESBL-Ec) is a global health concern that disproportionately affects sub-Saharan Africa (SSA). Gut mucosal colonisation is thought to precede invasive infection. Understanding ESBL-Ec colonisation and transmission across communities is therefore essential. We investigated the genomic epidemiology and spatial structure of 159 gut-colonising ESBL-Ec isolates from the faeces of 211 people in two rural Malawian villages using longitudinal sampling (2023–24), whole-genome sequencing and household mapping. Colonisation prevalence rose from 34.1% (95% CI: 27.8–41.0) to 54.2% (95% CI: 46.0–62.3) over one year. Isolates belonged to 33 sequence types (STs), most commonly ST38 and ST131, harbouring 46 distinct antimicrobial resistance gene types. Fifteen strains were identified in ≥3 households that were typically separated by short geographic distances (<400 m). Of 190 pairwise comparisons between same-strain isolates from different households sampled concurrently within villages, 88.9% differed by ≤10 single nucleotide polymorphisms, consistent with multi-household involvement in community transmission networks. Lineage-specific ST38 and ST131 network analyses linked rural isolates to urban Malawian isolates collected within the last decade. Our findings provide a transferable framework for inferring ESBL-Ec flow in community settings and highlight the need for One Health surveillance and improved sanitation infrastructure to limit transmission.

## Main Text

*Escherichia coli* is a species of gut commensal bacteria that can cause a range of clinical infections, including urinary tract infections (UTIs) and bloodstream infections (BSIs) (1). Antimicrobial resistance (AMR) among *E. coli* is a major global health issue, with *E. coli* being linked to more AMR-associated deaths in 2019 than any other bacterial species (>600,000) (2). AMR in *E. coli* is often mediated by the production of extended-spectrum beta-lactamases (ESBLs), which hydrolyse a broad range of beta-lactam antibiotics, including third-generation cephalosporins (3GCs) (3), thereby compromising a key first-line empirical treatment option. Difficulties in the treatment of infections caused by ESBL-producing *E. coli* (ESBL-Ec) are often compounded by the concurrent carriage of other antimicrobial resistance genes (ARGs) that can be co-located on mobile genetic elements (MGEs) such as plasmids and transposons (4,5), limiting the effectiveness of alternative antimicrobial agents.

Gut mucosal colonisation with ESBL-Ec is thought to precede BSI (7), is associated with increased risk of developing other invasive infections (8) and has rapidly increased in prevalence since the turn of the century (6). Understanding the colonisation and transmission dynamics of ESBL-Ec strains is a crucial step towards reducing morbidity and mortality in sub-Saharan Africa (SSA), where access to World Health Organisation (WHO) Watch and Reserve antibiotics is limited and where BSIs caused by ESBL-Ec are associated with increasing mortality (9,10). The surveillance of ESBL-Ec in low- and middle-income countries (LMICs) requires not only faecal sampling, but also sampling across a range of non-clinical settings, as there is a growing body of evidence demonstrating that AMR is a One Health problem in LMICs (11). The recent Drivers of Resistance in Uganda and Malawi (DRUM) study showed that flux of AMR bacteria between ecological compartments (humans, animals and the environment) enables ESBL-Ec transmission across eastern African communities, where water, sanitation and hygiene (WASH) infrastructure is often inadequate (12,13). A separate study in Blantyre (Malawi’s second largest city) showed that the diversity of healthcare-associated and community-acquired isolates was largely indistinguishable (14). Despite these knowledge advances, spatially resolved genomic analyses integrating household mapping with bacterial genomics have scarcely been conducted in rural SSA, leaving the dynamics of ESBL-Ec transmission and persistence within communities poorly understood in this context.

The resolution provided by whole-genome sequencing enables construction of ESBL-Ec transmission networks, but there is no standardised bioinformatic approach for inferring transmission events in communities. Various thresholds have been proposed to identify clonal complexes based on single nucleotide polymorphism (SNP) counts in core genome alignments. These range from strict thresholds (e.g., ≤5–10 SNPs) in outbreak studies (15) to loose thresholds (e.g., ≤100 SNPs) in One Health studies that investigate the flow of microbes between humans, animals and the environment (16). However, additional factors including the choice of reference genome, the relatedness of query genomes and the approach to handling genetic recombination can all affect core genome alignment lengths and resulting SNP counts (17), hence contextual interpretation is essential, as different methodological choices can be optimally suited to addressing different biological or epidemiological questions.

Here, we report a community-based genomic analysis of gut-colonising ESBL-Ec, integrated within a longitudinal cohort study originally established to investigate schistosomiasis transmission in two sentinel rural Malawian villages. We determine a rise in prevalence of gut colonisation after one year and identify inter-household strain-sharing networks. To contextualise geographically and temporally widespread lineages found in this study and in recent urban community- and hospital-derived isolate collections from Blantyre, we also construct sequence type (ST)-specific networks that assess the impact of recombination on core genome evolution, thereby enabling lineage-level inference of flow across urban and rural settings.

## Results

### Gut colonisation with ESBL-producing *E. coli* is dynamic and rising in prevalence

A total of 211 participants, aged 2–47 years, were recruited from 134 households. There were significant baseline differences in age and sex between recruits from Mangochi District and Nsanje District (*Table 1*). Participants in Nsanje District were generally older and more likely to be female than those recruited in Mangochi District. However, no significant differences in attrition rate, toilet facilities or drinking water source were detected between locations. Overall, 13.3% of participants reported that they did not use a latrine for defaecation. Most participants (96.7%) reported that they used a borehole for drinking water, with under 3% reporting use of an unprotected well or freshwater source.

**Table 1.** Baseline participant characteristics. Age (in years) is presented as median (IQR). Categorical variables are presented as proportions. *P-*values were calculated to assess differences in sample characteristics between study locations, using a Wilcoxon rank-sum test for age (numerical variable) and two-sided Fisher’s exact tests for categorical variables.

|  | Samama,<br>Mangochi District<br>(n = 163) | Mthawira,<br>Nsanje District<br>(n = 48) | P-value | Total<br>(n = 211) |
| --- | --- | --- | --- | --- |
| Demographics |  |  |  |  |
| Age (years) | 6 (4.0–12.0) | 14 (7.0–26.3) | <0.001 | 6 (4.0–16.5) |
| Female | 52.1 (85/163) | 77.1 (37/48) | 0.003 | 57.8 (122/211) |
| Study participation |  |  |  |  |
| Retained for follow-up | 75.5 (123/163) | 62.5 (30/48) | 0.097 | 72.5 (153/211) |
| Access to latrine for defaecation |  |  |  |  |
| Yes | 85.9 (140/163) | 87.5 (42/48) | 1.000 | 86.3 (182/211) |
| No | 13.5 (20/163) | 12.5 (6/48) |  | 13.3 (28/211) |
| Missing | 0.6 (1/163) | 0.0 (0/48) |  | 0.5 (1/211) |
| Drinking water source |  |  |  |  |
| Borehole | 95.7 (156/163) | 100.0 (48/48) | 0.490 | 96.7 (204/211) |
| Unprotected well or<br>freshwater source | 3.7 (6/163) | 0.0 (0/48) |  | 2.8 (6/211) |
| Missing | 0.6 (1/163) | 0.0 (0/48) |  | 0.5 (1/211) |

Among all 211 participants, baseline ESBL-Ec gut colonisation prevalence in late June–early July 2023 was 34.1% (95% CI: 27.8–41.0), as determined by isolation of ESBL-Ec from faeces (18) and downstream identification of ESBL genes in short-read draft genome assemblies using the ResFinder database (19). Among the 153 participants retained for 12-month follow-up in late June–early July 2024, prevalence rose from 33.3% (95% CI: 26.1–41.5) at baseline to 54.2% (95% CI: 46.0–62.3) at follow-up (McNemar’s test: *P* < 0.001). Of these participants, 28.1% (43/153) were not colonised with ESBL-Ec at either sampling point, 38.6% (59/153) became colonised, 17.6% (27/153) lost colonisation and 15.7% (24/153) were colonised at both sampling points (*Supplementary Fig. 1*). No significant difference in colonisation prevalence was detected between locations at baseline (Mangochi District = 34.4% [95% CI: 27.2–42.2], Nsanje District = 33.3% [95% CI: 20.8–48.5]; Chi-squared test: *P* = 0.896) or follow-up (Mangochi District = 52.8% [95% CI: 43.7–61.8], Nsanje District = 60.0% [95% CI: 40.8–76.8]; Chi-squared test: *P* = 0.481).

In total, 159 ESBL-Ec were isolated and sequenced, including 76 collected in 2023 and 83 collected in 2024. Based on a same-strain average nucleotide identity (ANI) threshold of 99.99% (20), the 159 isolates belonged to 83 different strains (*Supplementary Fig. 2*). In total, 62.3% (99/159) of the isolates belonged to a strain identified in ≥2 households, while 47.2% (75/159) belonged to one of 15 strains that were identified in ≥3 households. Isolates contained various plasmid replicon types, most commonly IncFIB (n = 115), IncFII (n = 66) and IncFIA (n = 50) (*Supplementary Fig. 3*). All isolates contained one *bla*_CTX-M_ ESBL gene, with *bla*_CTX-M-15_ being predominant (72.3%; 115/159) over both years (*Fig. 1a, Supplementary Fig. 4*). A total of 40 non-ESBL ARGs predicted to confer resistance against aminoglycosides, beta-lactams, chloramphenicol, fosfomycin, macrolides, quinolones, rifampicin, sulfonamides, tetracyclines or trimethoprim were detected. No carbapenemase genes were detected. ST profiles were dynamic between years (*Supplementary Fig. 5*), with a statistically significant difference detected in overall ST distributions between 2023 and 2024 (two-sided Fisher’s exact test with Monte Carlo simulation [10,000 replicates]: *P* < 0.001). ST38 (n = 23) and ST131 (n = 19) were the most prevalent among 33 distinct *E. coli* STs (*Fig. 1b*). A total of 29 popPUNK (Population Partitioning Using Nucleotide K-mers) lineages were identified (21,22). Two STs (ST10 and ST131) were associated with multiple popPUNK lineages each, while five popPUNK lineages included genomes from multiple STs. Of the 24 participants colonised both at baseline and at follow-up, in only one did we detect an isolate pair from the same ST and popPUNK lineage at both visits (ST131, popPUNK25), although the ANI between the genomes of the two isolates was 99.72% and 109 core genome SNPs were identified, indicating that the baseline isolate was likely not the ancestor of the follow-up isolate. In total, five *E. coli* phylogroups (A, B1, B2, D and F) were represented within the isolate collection, all of which were identified in both districts over the course of the study (*Fig. 1c*).

**Figure 1.**
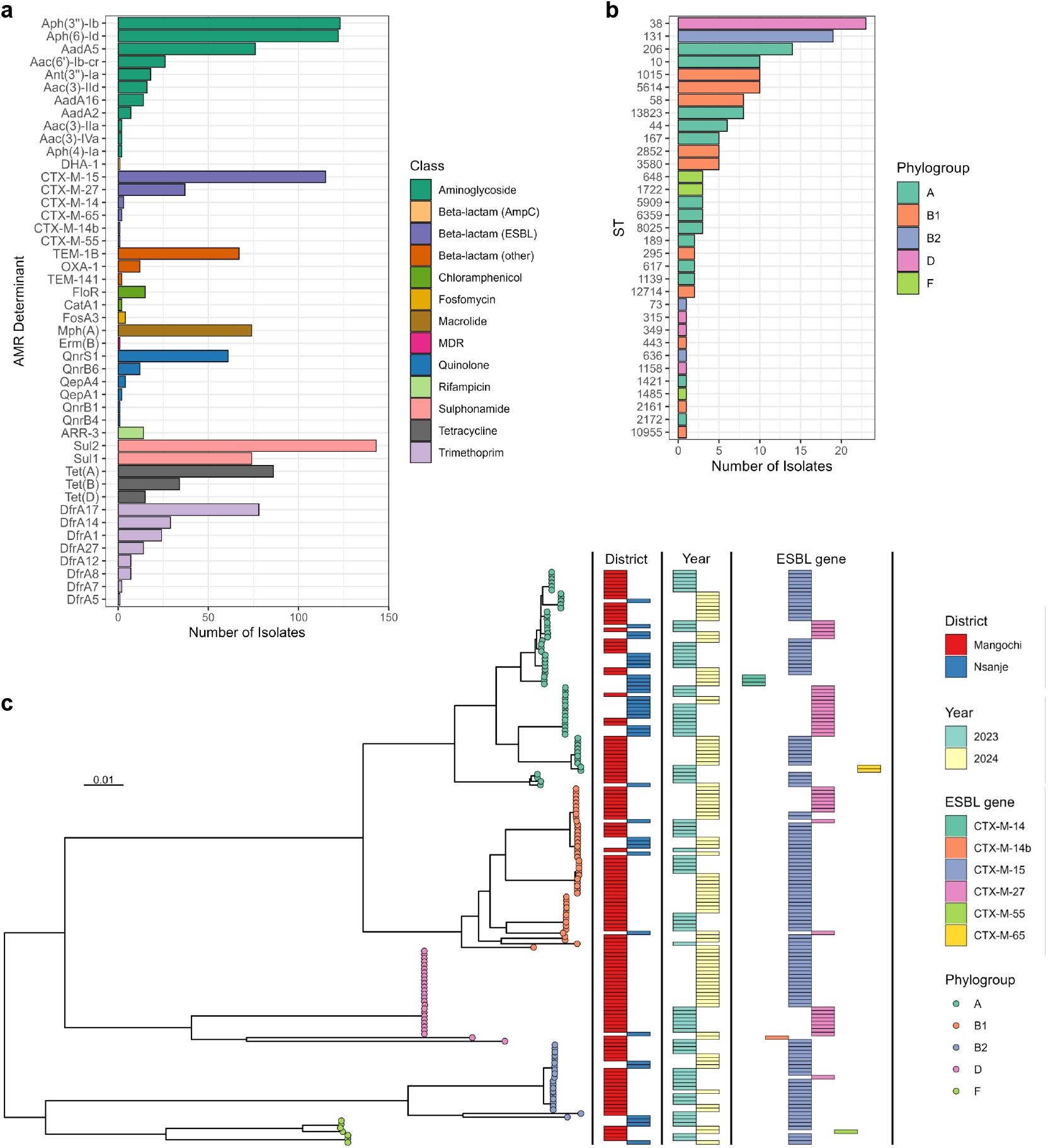
AMR determinants and phylogeny of ESBL-Ec isolated from faeces in rural Malawi. (a) Distribution of acquired AMR determinants. (b) Population structure by ST and phylogroup. (c) Midpoint rooted maximum likelihood phylogenetic tree. The tree scale bar indicates substitutions per site. The phylogeny illustrates the genetic diversity of ESBL-Ec circulating in rural Malawi, including five phylogroups shared across districts and years.

### Successful strains exhibit extensive ARG and virulence factor co-occurrence

Within the 15 strains identified in ≥3 households, from which a representative (medoid) isolate underwent long-read sequencing and hybrid assembly, we observed high levels of genome- and replicon-level ARG co-occurrence (median ARG count per genome = 9). ESBL genes (*bla*_CTX-M-15_, n = 12; *bla*_CTX-M-27_, n = 3) were mostly plasmid-borne (9/15). On these ESBL gene-harbouring plasmids of varying sizes (68.4–173.8 Kb) and replicon types (5 IncF, 3 IncY and 1 IncB/O/K/Z), 2–11 other (non-ESBL) ARGs were co-located *(Supplementary Table 1*). The incidence of clinical infection following gut colonisation was not investigated in this study. However, we assessed putative virulence by screening genomes against the Virulence Factor Database (VFDB). Genes spanning each of the major extraintestinal pathogenic *E. coli* virulence factor categories (adhesins, toxins, iron-uptake, protectins) were detected in all 15 strains found in ≥3 households. We also identified the *ybt* biosynthetic gene cluster (BGC) in 7/15 of these strains (*Supplementary Fig. 6*). This 11 gene yersiniabactin siderophore system promotes iron acquisition and has recently been associated with translocation of ESBL-Ec between the gut and the blood in neonates with bacteraemia in eastern Africa (23). Across all 159 ESBL-Ec genomes in our study, including those that were not distributed across multiple households, 196 distinct genes in the VFDB were identified. Diarrhoeagenic virulence genes that are major diagnostic targets for enteropathogenic *E. coli* (EPEC) and enterotoxigenic *E. coli* (ETEC) were identified (*eae* in 9 isolates from 5 strains, and *elt* in 1 isolate, respectively) (24). We did not detect Shiga toxin genes in any isolates.

### Strains form spatially structured networks involving nearby households

We investigated putative inter-household networks amongst the 15 strains identified in ≥3 households using strain-specific SNP calling. The 15 strains belonged to 11 STs from four phylogroups and were ordered for plotting based on descending maximum pairwise chromosomal SNP distance within each phylogroup (*Fig. 2a*). Three of these strains from ST10, ST206 and ST1015 were found in both districts. No isolate shared ≤10 chromosomal SNPs with any other isolate from a different district or year. Six of the 215 pairwise SNP distance comparisons within strains were between isolates from the same household, sharing between 0–2 SNPs. Most pairwise within-strain comparisons (88.4%; 190/215) were between isolates from the same village at the same sampling timepoint, but from different households. The mean pairwise SNP distance in these 190 comparisons was 3.90 (median = 2, range = 0–38). Most of the isolate pairs (88.9%; 169/190) differed by ≤10 SNPs and a further six pairs differed by 11–25 SNPs. In all 15 strains, a connection between isolates from different households differing by ≤10 SNPs was identified.

**Figure 2.**
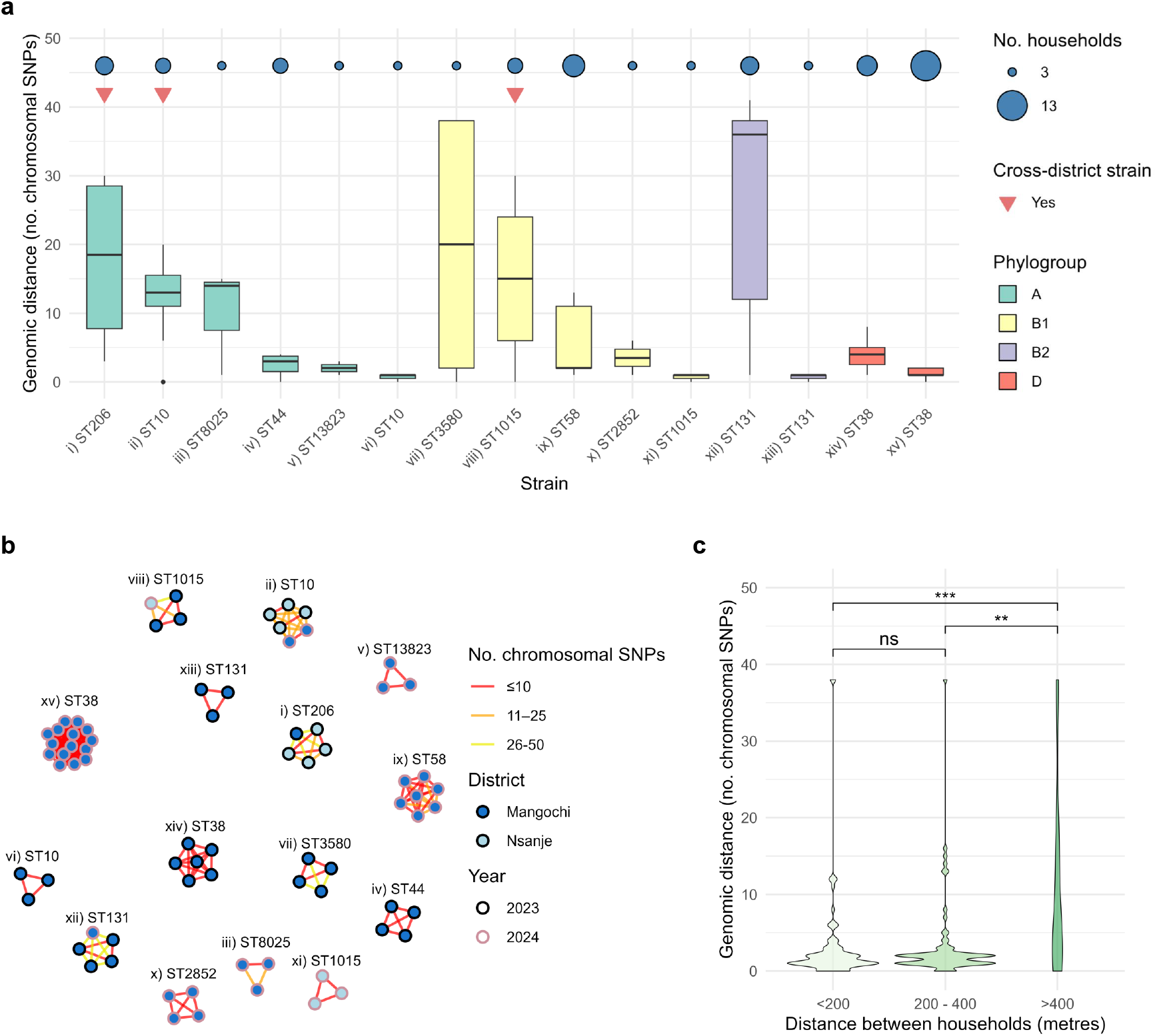
Genomic relatedness of ESBL-Ec isolates from strains identified in ≥3 households. (a) Boxplots showing the distribution of pairwise chromosomal SNP distances within each strain. Strains are ordered by descending maximum pairwise SNP distance within each phylogroup. Blue circles indicate the number of households contributing isolates to each strain (circle size proportional to household count) and red triangles indicate strains detected in both districts (cross-district strains). The centre lines indicate the median, boxes represent the interquartile range (IQR), and whiskers extend to the most extreme values within 1.5 × IQR. (b) SNP distance networks for each strain. Nodes represent individual isolates and edges represent pairwise SNP distances, coloured by distance category (≤10, 11–25 or 26–50 chromosomal SNPs). Node fill colour indicates location and outline colour indicates year of isolation. Networks illustrate close genetic relatedness of isolates within households and villages, with tighter clustering typically observed within the same district and sampling timepoint. (c) Violin plot showing the distribution of pairwise chromosomal SNP distances between ESBL-Ec isolates at the same sampling point and within the same district, but from different households, stratified by straight-line distance between households (<200 m, 200–400 m, and >400 m). Statistical significance was assessed using a Kruskal-Wallis test followed by two-sided Dunn’s post-hoc tests with correction for multiple comparisons; ns, non-significant; \**P* < 0.05 \*\**P* < 0.01; \*\*\**P* < 0.001.

Within the 15 strains, genomic and spatial distances were associated in pairwise comparisons between isolates sampled concurrently from different households in the same village. Those from hosts living >400 m apart tended to have higher SNP distances than pairs from hosts living <400 m apart (*Fig. 2c, Supplementary Fig. 7*). A Kruskal-Wallis test detected a statistically significant association (*P* = 0.001) between SNP distance and geographic distance category (<200 m, 200–400 m, >400 m). A post-hoc Dunn’s test did not detect a significant difference in the number of chromosomal SNPs between pairs from the <200 m and 200–400 m groups (*P* = 1.000), but the number of chromosomal SNPs was significantly greater in the >400 m group than both the <200 m group (*P* < 0.001) and the 200–400 m group (*P* = 0.008).

After anonymising household locations with a small random offset of ±50 m applied to both latitude and longitude, maps of strains identified in ≥3 households were plotted (*Fig. 3*). Within Samama, Mangochi District, where most faecal sampling was performed, some strains were clearly localised to one section of the village, including from strain xv (ST38) – represented by more isolates and households than any other strain (15 and 13, respectively). The other two strains involving >5 households in Samama – ix (ST58) and xiv (ST38) – also appeared to be both temporally and geographically constrained. Meanwhile, some strains were dispersed across the village, such as strain xii (ST131), that was also present at both sampling timepoints.

**Figure 3.**
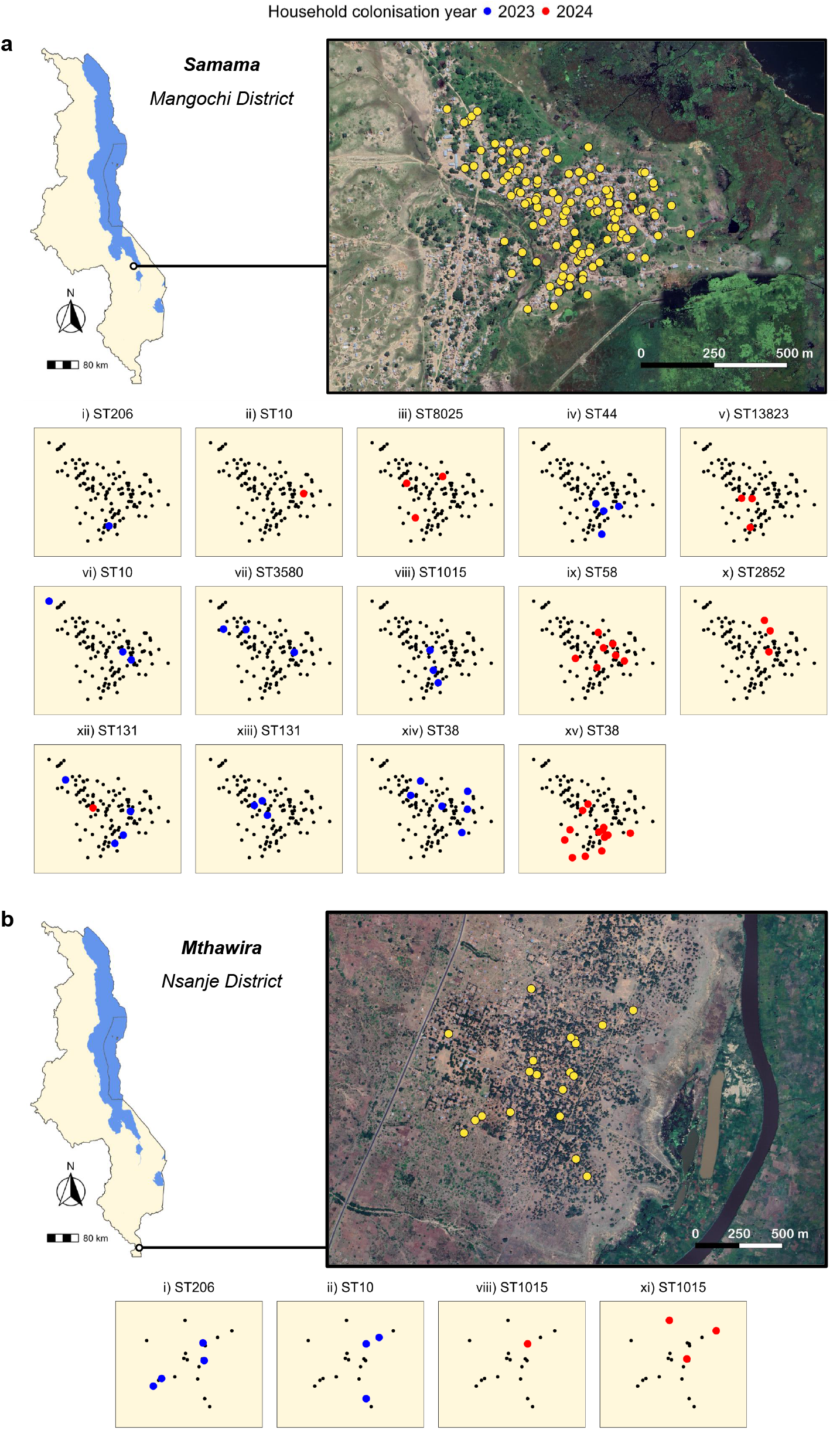
Spatial distribution of ESBL-Ec strains identified in ≥3 households. Maps show the distribution of strains in Samama, Mangochi District (a) and Mthawira, Nsanje District (b). For each strain, the underlying household map is shown, with all households represented as points. Larger coloured points are overlaid for households in which at least one participant was colonised with an isolate belonging to the corresponding strain (blue, 2023; red, 2024). To protect participant confidentiality, household GPS coordinates were anonymised by applying a small random offset (±50 m) to both latitude and longitude prior to plotting.

### Rural isolates from common STs are linked to urban and healthcare-associated isolates collected within the last decade

We constructed ST-specific phylogeny and networks for ST38 and ST131 isolates because these were the most abundant STs in our dataset and were represented across years and districts (23 and 19 isolates per ST, respectively). Both STs were also well represented in two faecal ESBL-Ec collections from Blantyre (the major city located between our two study sites), reported by Lewis et al., 2022 (isolates collected 2017–18) (14) and Musicha et al., 2025 (isolates collected 2019–20) (13). Lewis et al. sampled hospital inpatients, recently discharged patients and healthy community members, whereas Musicha et al. sampled exclusively from healthy community members. From these studies we identified and included 36 ST38 and 101 ST131 genomes, spanning healthy community members (25 ST38 and 66 ST131), recently discharged patients (9 ST38 and 32 ST131) and hospital inpatients (2 ST38 and 3 ST131), alongside genomes from rural community members collected in our study.

Prior to masking predicted recombinant regions, core genome alignments containing ATCG-only sites were 4,279,355 bp for ST38 and 3,954,219 bp for ST131. After masking predicted recombinant regions with Gubbins (25), alignment lengths reduced to 3,472,800 bp for ST38 and 2,834,919 bp for ST131. Of the polymorphic sites detected in the full alignments, 1,245/21,229 (5.86%) were retained in ST38 and 1,425/4,777 (29.83%) in ST131 after recombination masking. Thus, most polymorphisms in both STs were masked by Gubbins, and the number of polymorphic sites in core genome loci that were not affected by recombination was similar between ST38 and ST131. Despite this, the distribution of SNPs in recombination-masked core genome alignments differed markedly between STs. ST38 genomes were distributed across long-branch, deeply divergent clades (*Fig. 4a*), resulting in mean pairwise distances between isolates of different clades of almost 400 SNPs after regions impacted by recombination were masked (*Fig. 4c*). In contrast, branch lengths were shorter in the ST131 phylogeny (*Fig*. 4b) and SNP distances between clades were smaller (*Fig. 4d*). As a result, any two ST131 genomes differed at relatively few core genome sites once recombinant regions were excluded.

**Figure 4.**
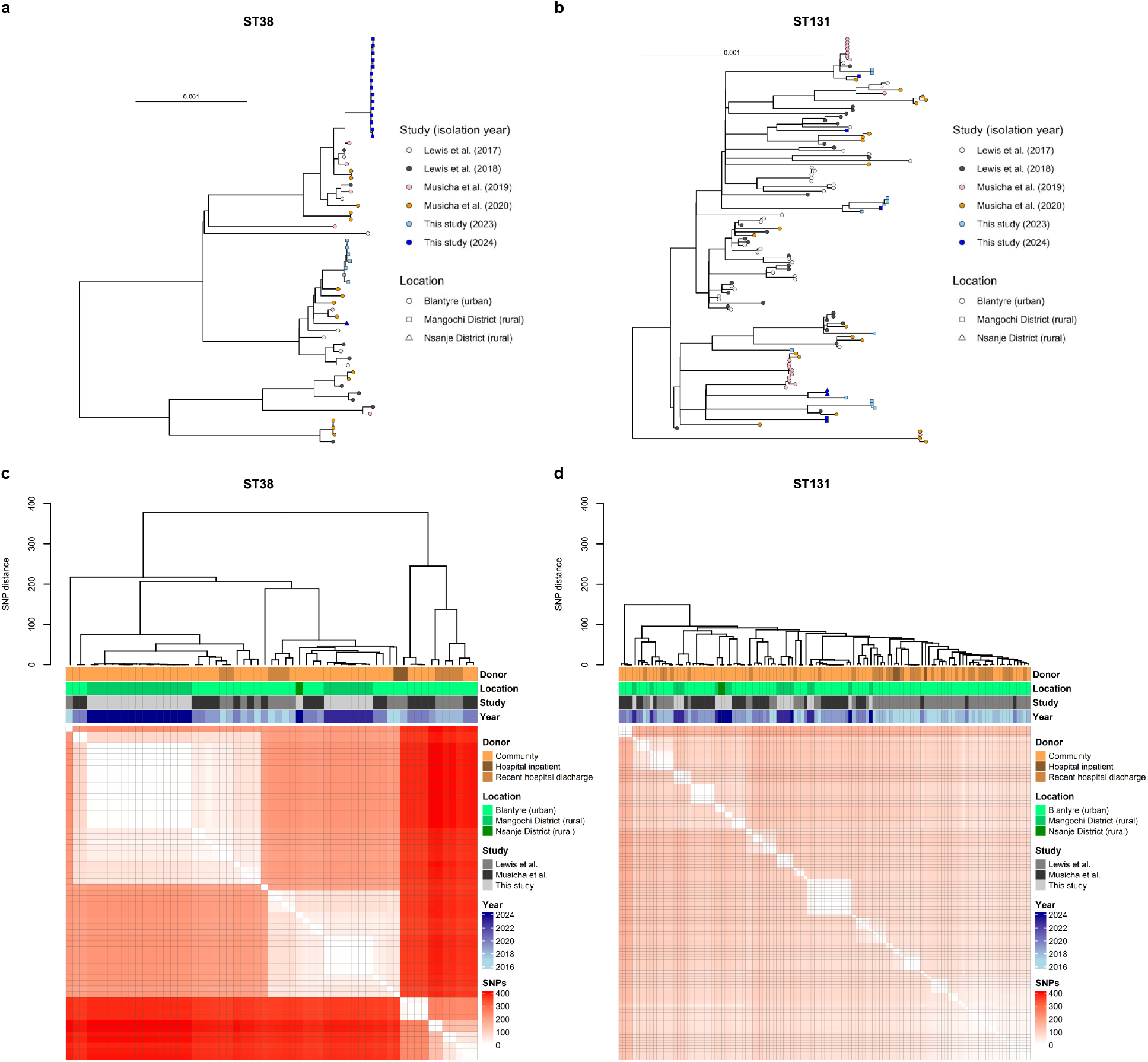
Midpoint-rooted maximum likelihood phylogenetic trees and recombination-masked chromosomal SNP distance heatmaps for major ESBL-Ec lineages circulating in rural and urban Malawi (ST38 and ST131). Genomes from this study are contextualised alongside ESBL-Ec genomes from previously published Malawian studies within the past decade, spanning rural (Mangochi District, Nsanje District) and urban (Blantyre) settings. In phylogenetic trees constructed for ST38 (a) and ST131 (b), tree tips indicate the study, year and location of the isolate genome. Tree scale bars indicate substitutions per site. SNP distance matrices with hierarchical clustering dendrograms are shown for ST38 (c) and ST131 (d). Column annotations indicate donor source (community, hospital inpatient, or recent hospital discharge), sampling location, contributing study, and year of isolation.

These lineage-specific evolutionary patterns were reflected in network plots. We constructed networks both with and without masking predicted recombinant regions to assess the impact of recombination masking on ESBL-Ec flow inference. For ST38, recombination masking had little effect on network structure (*Fig. 5a–b*). For ST131, however, masking recombinant regions substantially increased apparent connectivity among genomes (*Fig. 5c–d*). We implemented adjustments to arbitrary SNP distance thresholds when constructing networks to account for the reduction in core genome alignment lengths when masking predicted recombinant regions. *Escherichia coli* chromosomes are typically around 5 Mb long (22,26) and various arbitrary chromosomal SNP distance thresholds have been proposed for inference of clonal complexes. Ten SNPs across a 5 Mb alignment between two genomes (a strict threshold) is equivalent to variants at 0.0002% of sites, while 25 SNPs (a moderate and widely used threshold (17)) is equivalent to variants at 0.0005% of sites and 100 SNPs (a loose threshold) is equivalent to variants at 0.002% of sites. When applying these scaled thresholds to ST-specific networks, all rural ST38 genomes from our study (collected 2023–24) were linked to urban genomes from Lewis et al. and Musicha et al. (collected 2017–20) by loose links (25–99 SNPs per 5 Mb: <0.002%) both before and after removing recombinant regions. Variation ranging from <25 to >100 SNPs per 5 Mb (<0.0005% to >0.002%) was observed between the rural genomes and urban ST131 genomes both before and after masking recombination, but links were stronger and greater in number after masking recombination. This was despite adjusting transmission thresholds based on core genome alignment sizes (hence SNP counts were presented as proportions and count per 5 Mb). For example, for ST131, the maximum raw SNP count for inclusion in a moderate SNP-cluster (0.0005% SNP sites in the alignment, or 25 SNPs per 5 Mb) decreased from 19 to 14 after masking recombinant regions and adjusting the SNP threshold.

**Figure 5.**
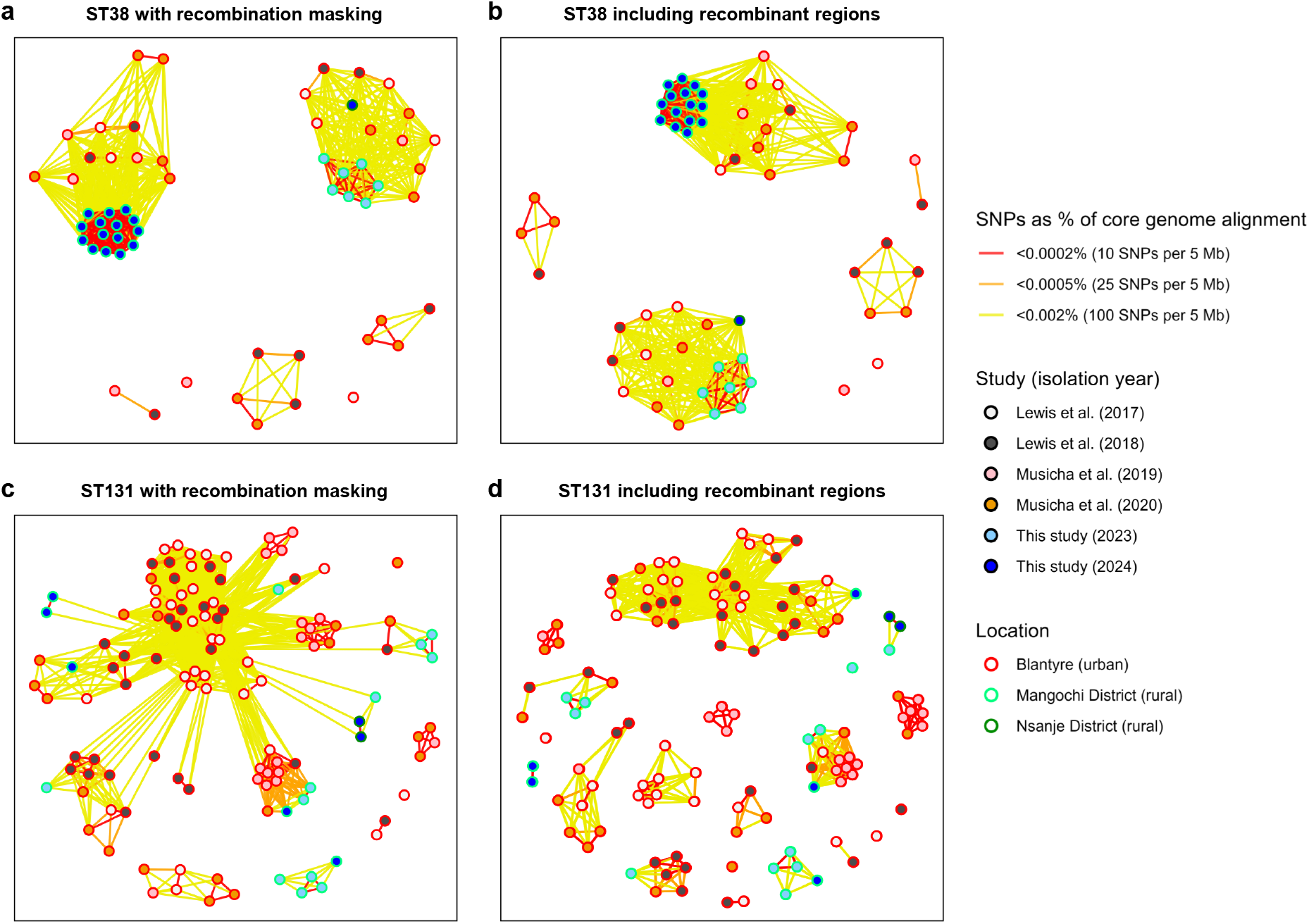
Core genome SNP distance networks for major ESBL-Ec lineages circulating in rural and urban Malawi (ST38 and ST131), with and without recombination masking. SNP-cluster networks are shown for ST38 (a,b) and ST131 (c,d) isolates, linking genomes from this rural study (Mangochi and Nsanje districts) with previously published urban isolates from Blantyre collected between 2017 and 2020. Networks were constructed both after masking recombinant regions (a,c) and using alignments including recombinant regions (b,d). Nodes represent individual genomes. Node fill colour indicates isolation year (and study). Node outline colour indicates sampling location. Edges connect genomes within defined pairwise SNP thresholds and are coloured by SNP distance as a percentage of the core genome alignment: <0.0002% (red; <10 SNPs per 5 Mb), <0.0005% (orange; <25 SNPs per 5 Mb), and <0.002% (yellow; <100 SNPs per 5 Mb).

Within the 2023 ST38 strict SNP-cluster (<10 SNPs per 5 Mb) identified in Mangochi District, one isolate (cS72c) belonged to a different ANI-defined strain to the other six, despite being highly similar across the core genome. The discordance was driven by differences in accessory genome content; although cS72c carried the same *bla*_CTX-M_ gene as the rest of genomes in the cluster, it possessed an additional *bla*_TEM_ gene, a different *dfrA* allele, and lacked several aminoglycoside and tetracycline ARGs present in the other isolates.

## Discussion

Understanding how ESBL-Ec circulate within and between communities is essential for designing strategies that limit their spread and prevent gut colonisation and subsequent clinical infection (13). In this 12-month follow-up study, we show that ESBL-Ec colonisation in rural Malawi is both highly prevalent and dynamic, frequently involving strain sharing between nearby households. We observed a substantial burden of ESBL-Ec gut colonisation, with prevalence exceeding 50% in the second year of the study. Colonisation was dynamic over the course of a year both in terms of the colonisation of individual hosts and the strains we identified (12). We did not identify evidence of ESBL-Ec carriage lasting the course of the study (one year) in any participants, as the isolates identified at baseline and follow-up belonged to different strains in participants colonised at both sampling points. However, it is likely that we underestimated diversity by taking a single colony pick of morphologically distinct phenotype. The flux observed in ST and strain composition contrasts with the persistent predominance of *bla*_CTX-M-15_ carriage within ESBL-Ec strains throughout the study. This particular ESBL gene is also widely prevalent across other eastern African locations (13,27), and globally (28).

Almost half of the isolates in our study belonged to one of 15 strains identified in ≥3 households. Co-occurrence of ARGs in whole genomes and co-localisation on plasmids amongst many of these successful strains underscores the potential clinical and public-health implications of strain and plasmid transmission, respectively, as does the presence of various virulence genes associated with extraintestinal infections. Should these ESBL-Ec strains cause BSIs or other invasive infections downstream (8), or should plasmids transfer to other pathogens (29), treatment options would be highly limited (9). Plasmids serve both as reservoirs and vectors of ARGs that can transfer across host bacteria and between environments in eastern Africa (13), but our findings suggest that limiting the spread of host lineages is also necessary to prevent widespread colonisation with ESBL-Ec.

The inter-household strain-sharing patterns identified in our study contrast with findings from a study of 50 households in urban Kenyan settlements, where *E. coli* strain-sharing was far more common within than between households (30). In that setting, human-to-human strain sharing among household members was strongly associated with contaminated drinking water (30). The near absence of strain sharing detected between households in said study may have been the result of the study’s sampling design, as only one household per urban compound was enrolled, preventing the detection of strain-sharing between households within the same compound. Differences in water-use behaviours and sanitation infrastructure between rural and urban areas may also affect how strains circulate between households (31,32). At baseline, 86.3% of participants in our study reported using latrines for defaecation, and 96.7% reported using a borehole for drinking water. As we did not sample latrine waste and drinking water, we were not able to incriminate WASH facilities in inter-household transmission. This remains a critical knowledge gap. Beyond water exposure, strain-sharing in human gut microbiomes is also shaped by social networks. A faecal shotgun metagenomic study conducted in 18 isolated villages in Honduras showed that individuals who are socially central often harbour strains representative of the broader village microbiome, and strain sharing can extend to non-familial and second-degree social connections (33). Together with these findings, our results indicate that ESBL-Ec transmission may occur through mixed pathways, potentially via contaminated water and interpersonal routes.

The structure of strain networks provides insight into the timescales of ESBL-Ec flow. Almost 90% of same-strain isolate comparisons from different households sampled concurrently within villages differed by ≤10 SNPs, consistent with recent transmission either directly between individuals or from a shared source, assuming typical *E. coli* mutation rates of <6 substitutions per genome per year in natural environments such as the gut (34–36). Distances between same-strain isolates in the range of 10–50 SNPs therefore likely represent shared ancestry more than a year in the past. The coexistence of these patterns suggests that the strains circulating between households are drawn from a pool of environmentally or socially maintained reservoirs. In combination with the transience of ESBL-Ec colonisation, these patterns point to a dynamic model of ESBL-Ec ecology in which strains and lineages repeatedly enter and exit human hosts, facilitated by both recent transmission events and longer-lived environmental reservoirs. Such complexity highlights why interventions targeting only household-level hygiene or antibiotic exposure are unlikely to fully interrupt the circulation of ESBL-Ec in these settings.

To explore links between isolates from ESBL-Ec lineages that are common in Malawi, we constructed ST-specific networks with and without masking predicted recombinant regions in core genome alignments. Masking recombination is recommended when studying recent evolutionary history (25), but it can hide meaningful genome dissimilarity introduced by horizontal gene transfer (HGT) and inflate the number of closely related isolate pairs (17). For this reason, Gorrie et al. argue that recombination should not routinely be masked when tracking the transmission of multidrug-resistant organisms across hospital networks, but they recognise that this approach, along with a single SNP threshold (25 in their case), may not be appropriate in all situations (17). One such situation may arise when multiple closely related strains from the same lineage are analysed, as recombination can generate divergent descendants from a recent common ancestor (37). Gorrie et al. also show that the effect of recombination masking on transmission inference can vary between STs (17). Our analyses across multiple community settings support this, as we identify contrasting evolutionary dynamics between two widespread Malawian ESBL-Ec STs. Most ST38 polymorphisms mapped to deep branches, consistent with older divergence. Consequently, transmission networks were largely insensitive to whether recombinant regions were included, yet genomes from rural and urban areas were loosely connected in the two predominant clusters by <100 SNPs per 5 Mb. In contrast, ST131 diversity collapsed into more connected clusters once recombination was masked, despite adjusting (reducing) SNP thresholds to account for the reduction in core genome alignment length. A greater number of ST131 genomes in our study were linked to urban genomes from 2017–20 by <25 SNPs per 5 Mb after this step. Recombination had therefore obscured some connectivity between rural and urban ST131 genomes. These findings underscore how recombination can distort phylogenetic signal, and potentially transmission inference, especially in lineages such as ST131 with higher than average recombination rates (38). We therefore suggest that transmission investigation across locations, hosts and environments should utilise both approaches (analyses with and without recombination masking) as the former may better reflect time to common ancestor (25) while the latter provides greater resolution across the full core genome (17).

Our results also highlight an important conceptual distinction between ANI-based strain identification and core genome SNP-based transmission network analyses. With a same-strain threshold of >99.99% ANI, genomes typically share high gene-content similarity (>99.0%) (20,39), therefore sharing key phenotypic traits such as those related to AMR (20). Although genomes from the same strain may also belong to the same clonal complex, supporting evidence from lineage-specific SNP analyses are needed to infer clonality and transmission. In principle, two ESBL-Ec genomes with 500 SNPs over a 5 Mb core genome alignment can share 99.99% ANI. However, our data show that this is very unlikely to occur in nature, as HGT can rapidly introduce accessory genome divergence between clonal isolates with near identical core genomes. No isolate pairs in our study shared >99.99 ANI and ≥50 chromosomal SNPs. Meanwhile, the ST38 core genome alignment showed that one ST38 genome in our study shared <10 SNPs per 5 Mb with the other six ST38 genomes also isolated in Mangochi District in 2023, but this genome formed its own ANI-defined strain due to differences in accessory genome content, including ARGs. This example illustrates how rapid acquisition, loss and exchange of MGEs can cause isolates with recent common ancestry to diverge into different ANI-defined strains, whereas core genome SNPs more reliably capture their underlying evolutionary relatedness.

These findings inform a set of practical recommendations for inferring ESBL-Ec flow within and between communities. Firstly, same-strain genomes sharing >99.99% ANI and exhibiting high similarity across core and accessory components are likely to share a recent common ancestor when supported by epidemiological data. However, SNP analysis remains essential for transmission network construction, with >10 SNPs likely indicating >1 year since a shared common ancestor. We propose constructing transmission networks both with and without masking predicted recombinant regions when analysing isolates from the same lineage but different strains, to account for the confounding effects of HGT in the former while preserving informative dissimilarity signal in the latter. We further recommend adjusting SNP-based transmission thresholds to account for core genome alignment length reduction, particularly when masking variation in regions affected by recombination. Here, 0.0002% SNPs in genome alignment equates to approximately 10 substitutions across a 5 Mb *E. coli* genome, which we observed between genomes collected in the same year, whereas links across years and locations may more commonly be identified using moderate to loose clustering thresholds equivalent to <25 or <100 SNPs per 5 Mb in the core genome (<0.0005% and <0.002%, respectively).

Our study has several limitations. Our analysis focused on *E. coli* isolates cultured on selective agar, therefore excluding non-ESBL-producing *E. coli* that also influence colonisation dynamics via competition in the gut microbiome (40). Additionally, we likely underestimated the diversity of gut-colonising ESBL-Ec and overestimated the transience of colonisation, as we only took a single colony pick from each morphologically distinct phenotype due to resource constraints. Although we captured substantial temporal dynamics using two sampling timepoints, additional timepoints would have helped us to characterise colonisation duration. Our identification of strains circulating across districts would have benefited from denser sampling in Nsanje District, where fewer participants were recruited. However, this study was embedded within a pre-existing longitudinal schistosomiasis surveillance programme in southern Malawi, and only participants providing faecal samples within that framework were eligible for inclusion. While urogenital schistosomiasis is endemic across much of Malawi, intestinal schistosomiasis occurs more focally, including in Mangochi District, and predominantly affects children (41). Consequently, a large proportion of samples analysed in this study were obtained from children in Mangochi District. Finally, while we did detect strong inter-household connectivity, we did not undertake parallel sampling of waste systems and drinking water, nor did we sample stagnant water bodies formed through flooding, which are increasingly common in the villages and may contribute to the translocation of ARGs (12,42,43), preventing us from formally describing transmission chains. These limitations highlight the need for integrated, multi-compartment, longitudinal sampling to better understand ESBL-Ec transmission pathways in rural Malawi (11,44).

In summary, our data demonstrate that gut colonisation with ESBL-Ec in rural Malawi is highly prevalent, and that successful strains are shared between members of nearby households. Integrating waste system, drinking water, environmental and livestock sampling alongside human sampling will be essential to monitor clonal expansion and identify transmission pathways in and between communities. Effective control of ESBL-Ec transmission in communities is likely to require strategies that extend beyond single households or clinical settings, including improvements in WASH infrastructure.

## Methods

### Study design and data handling

The bacterial isolates described here were collected in collaboration with a pre-established longitudinal freshwater parasite surveillance study (Hybridisation in Urogenital Schistosomiasis; HUGS) (42) in two sentinel villages in rural southern Malawi: Samama, Mangochi District (S 14.418767°, E 35.220985°), and Mthawira, Nsanje District (S 16.849802°, E 35.290041°) (41,45). Within the HUGS study, all participants provided annual urine samples for urogenital schistosomiasis diagnostics, but the provision of faecal samples for intestinal schistosomiasis diagnostics was optional as the tracking of intestinal parasites was not a primary objective. In this AMR study, we sought to maximise the utility of faecal samples by gaining consent for samples to undergo microbiological analysis in addition to routine parasite diagnostics (44). Participants aged 2 years or older providing faecal samples within the HUGS study were eligible for inclusion in this AMR study, provided they were not acutely unwell at the time of sampling and additional consent was acquired for their samples to undergo microbiological analysis. Participants under the age of 16 required additional consent from a parent or guardian. Ethical approvals were granted by The College of Medicine Research Ethics Committee, Malawi (approval no. P.06/23/4135) and the Liverpool School of Tropical Medicine Research Ethics Committee, United Kingdom (approval no. 22-028).

In late June–early July 2023, 211 people aged 2–47 years providing faecal samples within the HUGS study were consented for inclusion in this study, including 163 participants from 114 households in Samama (Mangochi District) and 48 participants from 20 households in Mthawira (Nsanje District). Baseline participant characteristics, including demographic and WASH data, were collected via questionnaires. We attempted to obtain repeat faecal samples from all participants 12 months later. 72.6% (153/211) of participants provided follow-up faecal samples in late June–early July 2024, including 123 from 90 households in Samama and 30 from 14 households in Mthawira.

### Sample processing and data handling

Faecal sample processing followed optimized methods for the targeted surveillance of ESBL-Ec in human faeces (18). Briefly, bacteria from faecal swabs underwent a 4-hour pre-enrichment in Buffered Peptone Water, before 1 µL of pre-enrichment media was plated out on ESBL-selective chromogenic agar (CHROMagar ESBL, CHROMagar). A single colony pick of each morphological phenotype was taken, before species was confirmed using a MALDI-TOF mass spectrometer (Bruker). ESBL production was confirmed using the combination disc method on Mueller-Hinton agar with discs of cefotaxime (30 μg) and ceftazidime (30 μg) with 10 μg clavulanic acid (Becton Dickinson) and without clavulanic acid (Oxoid), in accordance with UK standards for Microbiology Investigations (46). ESBL production was confirmed if there was a difference of 5 mm or more between the clavulanic acid and non-clavulanic acid discs for either cephalosporin (14,46).

Participant and isolate data were imported into the R Studio environment (R Version 4.5.2) for downstream analysis. To assess differences in sample characteristics between study locations, a Wilcoxon rank-sum test was used for age (numerical variable) and two-sided Fisher’s exact tests were used for categorical variables. A McNemar’s test was used to check for a significant change in ESBL-Ec gut colonisation prevalence amongst participants providing baseline and 12-month follow-up samples in 2023 and 2024. Prevalence estimates were calculated as proportions with 95% confidence intervals (CIs) using the Wilson score method with continuity correction, implemented via R’s ‘prop.test’ function. Chi-squared tests were used to assess for differences in prevalence between districts at both sampling timepoints.

### Short-read sequencing and processing

All ESBL-Ec isolates were shipped to MicrobesNG (UK) for DNA extraction, library preparation and Illumina short-read sequencing with the Illumina NovaSeq 6000 instrument using a 2×250 bp paired-end configuration. The next steps of processing were conducted upon delivered FASTQ sequence files at the Liverpool School of Tropical Medicine (LSTM). Raw FASTQ files were quality assessed using FASTQC (https://github.com/s-andrews/FastQC) (Version 0.11.9). Trimming with Trimmomatic (47) (Version 0.39) was performed to remove adapter sequences and leading or trailing bases with a quality score <4, bases with a mean quality score <20 over a sliding window of four bases, and any reads with length below 36 following removal of low-quality bases. De novo assemblies were generated using the Shovill pipeline (https://github.com/tseemann/shovill) (Version 1.0.4) with SPAdes (48) (Version 3.13.0), specifying an expected genome size of 5 Mb, an input read coverage of ∼100×, and a minimum contig length of 500 bp. Assembly statistics were calculated with QUAST (49) (Version 5.0.2) and CheckM (50) (Version 1.1.2). All assemblies were >4 Mb in length (range = 4.55–6.05), completeness ranged from 99.07–100.00% and contamination ranged from 0.04–1.14%. All draft genomes were therefore deemed high-quality (51) and were included in downstream analysis. Using ABRicate (https://github.com/tseemann/abricate) (Version 1.0.1), assemblies were screened against the ResFinder database (19), the Plasmidfinder database (52) and the VFDB (53) (all downloaded 11 February 2025) to identify ARGs, plasmid replicons and virulence genes, respectively, specifying a minimum nucleotide identity of 90% and a minimum coverage of 80% to call a hit.

### Phylogenetics

STs were determined using MLST (https://github.com/tseemann/mlst) (Version 2.23.0) with the seven-gene Achtman scheme. To test for change in ST distribution dynamics between years, a two-sided Fisher’s exact test was performed with Monte Carlo simulations (10,000 replicates) due to sparse counts for many STs in the dataset. Annotation- and alignment-free population clustering was performed with popPUNK (21) (Version 2.7.5) using the *E. coli* v2 database of globally representative *E. coli* genomes (22) as reference. Next, fastANI (54) (Version 1.3.4) was used to perform pairwise ANI calculations and generate an ANI matrix. Hierarchical clustering was performed on a distance matrix (1 - (ANI / 100)) in the R environment using the base R ‘hclust’ function with average linkage, where the distance between two clusters is defined as the average pairwise distance between all isolates in those clusters. Strain groups were defined as clusters in which the average pairwise ANI among members exceeded 99.99% (20,39).

To construct phylogeny, a map-to-reference core genome alignment was performed using the Snippy pipeline (https://github.com/tseemann/snippy) (Version 4.6.0) at default settings, with the *E. coli* K-12 genome as reference (RefSeq accession: GCF_000005845.2), before Gubbins (25) (Version 2.3.4) was used to identify and mask hypervariable areas of the genome, representing likely recombinant regions. Polymorphic sites between the ESBL-Ec isolates in our study were then extracted from the recombination-masked core genome alignment using SNP-sites (55) (Version 2.5.1). A maximum likelihood phylogenetic tree was then constructed with IQ-TREE (Version 1.6.1) using the ModelFinder option (TVM+F+ASC+G4 chosen) and 1,000 bootstraps. The tree was imported into R as a “phylo” object with the R/Bioconductor *ape* (Version 5.8.1) package, midpoint-rooted with the R/Bioconductor *phangorn* (Version 2.12.1) package, then prepared for downstream visualisation and annotation.

### Long-read sequencing and processing

Within each strain found in 3 or more households, we identified the medoid genome for long-read sequencing (the genome with the highest mean ANI compared to all other isolates in that strain). Medoid genomes represent the least-divergent genotype within a set of closely related genomes and are routinely used as representative genomes in large-scale bacterial taxonomy and genome catalogues (56–58). In the absence of known ancestral sequences, which are unlikely to have been captured by our sampling, this approach provided an estimate of the isolate closest to the common ancestral genotype within each strain. Medoid strains were shipped to Plasmidsaurus (USA), for DNA extraction, library preparation, Oxford Nanopore long-read sequencing using the R10.4.1 flow cell, base calling and adapter trimming. The next steps of processing were conducted upon receiving FASTQ sequence files at LSTM.

Filtlong (https://github.com/rrwick/Filtlong) (Version 0.2.1) was used to discard reads shorter than 1 Kb, after which the lowest-quality 10% of remaining reads were removed using default quality weighting. Consensus long-read assemblies were then performed using the Autocycler pipeline (59) (Version 0.5.2). Within the Autocycler environment, genome size was estimated with Autocycler helper functions, then four subsampled read sets were generated for each sample using the Autocycler subsampler function, with optimal coverage for each subset determined by the total read depth and estimated genome size to ensure sufficient read depth and independence of each read subset. Four assemblies were then performed upon each read subset before Autocycler generated an output consensus assembly for each genome. The four different long-read assemblers used within the Autocycler environment were Canu (60), Flye (61), NECAT (62) and Plassembler (63). Genomes were reoriented with Dnaapler (64) (Version 1.3.0), before three rounds of polishing were conducted to produce hybrid assembled genomes, first with long-reads using Medaka (https://github.com/nanoporetech/medaka) (Version 2.0.1), then with short-reads using Polypolish (65) (Version 0.5.0) and Pypolca (66) (Version 0.3.1) using the ‘--careful’ option. Genome assemblies were annotated using Bakta (67) (Version 1.10.4; full database Version 5.1). ARGs, plasmid replicons and virulence genes were identified by screening assemblies with ABRicate, as earlier described for short-read assemblies in our study.

Short-reads from non-medoid genomes within each strain were mapped to their respective hybrid assembled medoid chromosomal reference genomes before polymorphic sites were identified and extracted with SNP-sites, as earlier described. Pairwise SNP distances between genomes from the same strain were calculated using snp-dists (https://github.com/tseemann/snp-dists) (Version 0.8.2). SNP distance matrices for genomes within each strain were used to generate strain-specific SNP networks using the R/CRAN *igraph* (Version 2.2.1) package.

### Geospatial analysis

To enable linkage of genomic and geographic distances between isolates from the same strain sampled concurrently from different households within the same village, geographic distances between isolate hosts were calculated from household GPS coordinates using the R/CRAN *geosphere* (Version 1.5.20) package. A Kruskal-Wallis test was conducted to assess whether there was a statistically significant association between SNP distance and geographic distance category (<200 m, 200–400 m, >400 m) among same-strain genomes from different households. A post-hoc Dunn’s Test with Bonferroni correction was then performed using the R/CRAN *FSA* (Version 0.10.0) package. To protect participant anonymity while preserving the general household structure in maps, a small random offset of ±50 m was applied to both latitude and longitude before saving anonymized coordinates to the publicly available dataset that was used to plot jittered strain distribution between households.

### Urban-rural transmission network construction and phylogeny

To investigate the relatedness of rural faecal ESBL-Ec isolates sequenced in our study and ESBL-Ec isolates from urban Malawian faecal samples in two recently published studies (13,14), we downloaded the sequences of 139 ST38 and ST131 ESBL-*E. coli* genomes collected in Blantyre, Malawi from the European Nucleotide Archive (project IDs: PRJEB26677, PRJEB28522, PRJEB37378). The study of Musicha et al. included genomes from other eastern African locations (13), but only those from Blantyre – the urban centre between our two rural sampling sites – were included in our analysis. Downloaded sequences underwent quality control, read trimming, assembly and MLST confirmation using the same methods as for genomes sequenced in our study. Two genomes labelled as ST131 were excluded after MLST because of imperfect matches for the *gyrB* allele, which prevented confident ST assignment. A total of 137 genomes from these studies were therefore included in further analysis (ST38: n = 36; ST131: n = 101), alongside the ST38 and ST131 genomes sequenced in our study.

Before constructing core genome alignments, we retrieved external ST-specific references: *E. coli* strain NCTC 13441 (NCBI accession: NZ_LT632320.1) for ST131 and *E. coli* strain 114 (NCBI accession: NZ_CP023364.1) for ST38. Reads from each ST collection were mapped to their respective reference genome, before a core genome alignment for each ST was constructed, hypervariable (recombinant) regions were predicted, polymorphic sites were extracted and pairwise SNP distances were calculated, as earlier described. Polymorphic site extractions and pairwise SNP distance calculations were performed on core genome alignments both before and after masking predicted recombination sites with Gubbins. To assess the differences in core genome alignment sizes and polymorphic sites as a result of masking hypervariable regions, a recombination-masked alignment that included monomorphic sites was generated using the “mask_gubbins_aln.py” (https://github.com/nickjcroucher/gubbins/blob/masking_aln/python/scripts/mask_gubbins_aln.py) post-processing script (without this, Gubbins only outputs polymorphic sites), then both monomorphic and polymorphic ATCG-only sites across genomes were extracted using SNP-sites and alignment files were compared using SeqKit (68) (Version 2.1.0). ST-specific networks were constructed from SNP distance matrices using strict, moderate and loose cluster thresholds corresponding to 10, 25 and 100 SNPs per 5 Mb, respectively. Networks were generated from core genome alignments both before and after masking recombination. Phylogenetic trees were constructed for each ST as earlier described, using IQTREE upon alignments containing only polymorphic sites extracted from recombination masked core genomes. The TVMe+ASC and K3P+ASC models were selected for ST38 and ST131, respectively, by ModelFinder.

### Data visualisation

The Clinker webtool (69) (https://cagecat.bioinformatics.nl/tools/clinker) was used to visualise gene cluster comparisons between genomes containing the BGCs associated with translocation of ESBL-Ec from the gut to the blood. All other data visualisation was using base R functions and the R/CRAN *ggplot2* (Version 4.0.1), *ggpubr* (Version 0.6.2), *ggraph* (Version 2.2.2), *ggspatial* (Version 1.1.10) and *ComplexHeatmap* (Version 2.26.1) packages, as well as the R/Bioconductor *ggtree* (Version 4.0.4) and *ggtreeExtra* (Version 1.20.1) for visualisation of annotated phylogenetic data (70).

## Author statements

### Sequence and data availability

Short- and long-reads from isolates sequenced in this study have been deposited in the Sequence Read Archive (SRA) under BioProject PRJNA1399404. SRA accession numbers are SRR36707243– SRR36707401 for short-reads and SRR36746900–SRR36746914 for long-reads. Data and code used to carry out analyses in R are available at https://github.com/amoreo71/malawi_esbl_e_coli_genomics.

### Author contributions

Conceptualisation: AMO’F, JM, JRS and APR; Data curation: AMO’F, DL, PMa, GN, JML, PMu, SAK, JM, JRS and APR. Formal analysis: AMO’F, DL, JML, RG, EA, SM, JRS, APR; Resources: JM, JRS, APR; Writing – original draft: AMO’F; Writing – reviewing and editing: all authors.

### Conflicts of interest

None to declare.

### Funding

AMO’F is funded by the Medical Research Council via the Liverpool School of Tropical Medicine / Lancaster University doctoral training partnership (grant no. MR/W007037/1). Fieldwork was supported by the Wellcome Trust within a Joint Investigator Award (grant no. 220818/Z/20/Z). APR acknowledges funding from the Medical Research Council, Biotechnology and Biological Sciences Research Council and Natural Environmental Research Council which are all Councils of UK Research and Innovation (grant no. MR/W030578/1) under the umbrella of the JPIAMR (Joint Programming Initiative on Antimicrobial Resistance), and UKRI through the Strength in Places Fund (grant no. SIPF 36348). The MALDI-TOF was purchased using grant funding from the Medical Research Council (grant no. MC_PC_MR/Y002466/1). The funders had no role in study design, data collection and analysis, decision to publish, or preparation of the manuscript.

## Acknowledgements

We are grateful to all participants in the study. We thank field staff associated with the Hybridisation in Urogenital Schistosomiasis (HUGS) study from the Malawi Liverpool Wellcome Research Programme and the Liverpool School of Tropical medicine, including Alexandra Juhász, Sam Jones, Ruth Cowlishaw, Sarah Rollason, Priscilla Chammudzi, Donales R. Kapira, E. James LaCourse, Bright Mainga, Clinton Nkolokosa and John Archer. Additionally, we thank Simon Wagstaff, Andrew Bennet and Sophie Dunkley at LSTM for technical support.

## Notes

https://github.com/amoreo71/malawi_esbl_e_coli_genomics

### Competing Interest Statement

The authors have declared no competing interest.

