## Supplementary materials for "Whole-genome sequencing reveals inter-household networks of gut-colonising ESBL-producing *Escherichia coli* in two rural Malawian districts"

Supplementary Table 1. Supplementary Table 1. ARG co-occurrence and virulence in medoid genomes of strains identified in ≥3 households.

Supplementary Figure 1. ESBL-Ec gut colonisation groups after baseline and 12-month follow-up sampling.

Supplementary Figure 2: ANI hierarchical clustering dendrogram.

Supplementary Figure 3: Plasmid replicon types.

Supplementary Figure 4: ARG profile dynamics (2023–24).

Supplementary Figure 5: ST profile dynamics (2023–24).

Supplementary Figure 6: Yersiniabactin (ybt) biosynthetic gene cluster comparison.

Supplementary Figure 7: Scatter plot of pairwise SNP distances between same-strain isolates identified in ≥3 households.

**Supplementary Table 1. ARG co-occurrence and virulence in medoid genomes of strains identified in ≥3 households.** For each strain, the table reports the representative isolate, the ESBL gene detected and its genomic context, ARGs located on the same replicon as the ESBL gene and ARGs located on other replicons. Presence or absence of the yersiniabactin biosynthetic gene cluster (ybt BGC) and diarrhoeagenic pathotype are also shown.

| Phylogroup | Strain no. (ST) | Isolate | ESBL gene (associated replicon type) | ARGs on replicon with ESBL gene | ARGs on other replicons | *ybt* BGC | Diarrhoeagenic pathotype |
|---|---|---|---|---|---|---|---|
| A | i (ST206) | cM2a | *bla*$_{CTX-M-27}$ (IncFIB plasmid) | *aac(6')-Ib-cr, aadA5, aadA16, ARR-3, dfrA17, dfrA27, floR, qnrB6, sul1, sul2, tet(A)* | - | No | - |
| | ii (ST10) | cM8a | *bla*$_{CTX-M-15}$ (IncB/O/K/Z plasmid) | *dfrA1, qnrS1* | *aph(6)-Id, aph(3'')-Ib, sul2, tet(A)* | Yes | - |
| | iii (ST8025) | fS149a | *bla*$_{CTX-M-15}$ (chromosome) | *mph(A), tet(B)* | *aph(3'')-Ib, aph(6)-Id, bla*$_{TEM-1B}$*, dfrA8, mph(A), sul2* | Yes | - |
| | iv (ST44) | cS76a | *bla*$_{CTX-M-15}$ (IncFIA/IncFIB plasmid) | *aadA5, dfrA17, mph(A), sul1, tet(B)* | *aph(3'')-Ib, aph(6)-Id, sul2* | Yes | - |
| | v (ST13823) | fS25a | *bla*$_{CTX-M-15}$ (IncFIB plasmid) | *aadA5, dfrA17, qnrS1, sul1* | *aph(3'')-Ib, aph(6)-Id, sul2* | No | EPEC |
| | vi (ST10) | cS100a | *bla*$_{CTX-M-15}$ (chromosome) | - | *aac(3)-IId, bla*$_{TEM-1B}$ | Yes | - |
| B1 | vii (ST3580) | cS108a | *bla*$_{CTX-M-15}$ (chromosome) | *qnrS1* | *aph(3'')-Ib, aph(6)-Id, bla*$_{TEM-1B}$*, dfrA8, sul2* | No | - |
| | viii (ST1015) | cS33a | *bla*$_{CTX-M-15}$ (IncY plasmid) | *aph(3'')-Ib, aph(6)-Id, bla*$_{TEM-1B}$*, dfrA14, qnrS1, sul2* | *tet(B)* | No | - |
| | ix (ST58) | fS54a | *bla*$_{CTX-M-27}$ (IncFIA/IncFIB plasmid) | *aadA5, aph(3'')-Ib, aph(6)-Id, dfrA17, mph(A), sul1, sul2, tet(A)* | - | No | - |
| | x (ST2582) | fS58a | *bla*$_{CTX-M-15}$ (IncY plasmid) | *aph(3'')-Ib, aph(6)-Id, bla*$_{TEM-1B}$*, dfrA14, qnrS1, sul2, tet(A)* | *bla*$_{TEM-1B}$*, mph(A)* | No | - |
| | xi (ST1015) | fM51a | *bla*$_{CTX-M-15}$ (IncY plasmid) | *aph(3'')-Ib, aph(6)-Id, bla*$_{TEM-1B}$*, dfrA14, qnrS1, sul2* | *tet(B)* | No | - |
| B2 | xii (ST131) | cS53a | *bla*$_{CTX-M-15}$ (chromosome) | *aac(6')-Ib-cr, bla*$_{OXA-1}$ | *aac(3)-IId, aadA5, aph(3'')-Ib, aph(6)-Id, bla*$_{TEM-1B,}$ *dfrA17, mph(A), sul1, sul2, tet(A)* | Yes | - |
| | xiii (ST131) | cS91a | *bla*$_{CTX-M-15}$ (chromosome) | - | *aadA5, aph(3'')-Ib, aph(6)-Id, bla*$_{TEM-141}$*, dfrA17, mph(A), sul1, sul2, tet(A)* | No | - |
| D | xiv (ST38) | cS51a | *bla*$_{CTX-M-27}$ (IncFIB/IncFII plasmid) | *aadA5, aph(3'')-Ib, aph(6)-Id, dfrA17, mph(A), sul1, sul2, tet(A)* | - | Yes | - |
| | xv (ST38) | fS10a | *bla*$_{CTX-M-15}$ (chromosome) | *ant(3'')-Ia, aph(3'')-Ib, aph(6)-Id, bla*$_{TEM-1B}$*, dfrA1, sul2, tet(D)* | - | Yes | - |

**Supplementary Figure 1. ESBL-Ec gut colonisation groups after baseline and 12-month follow-up sampling.**

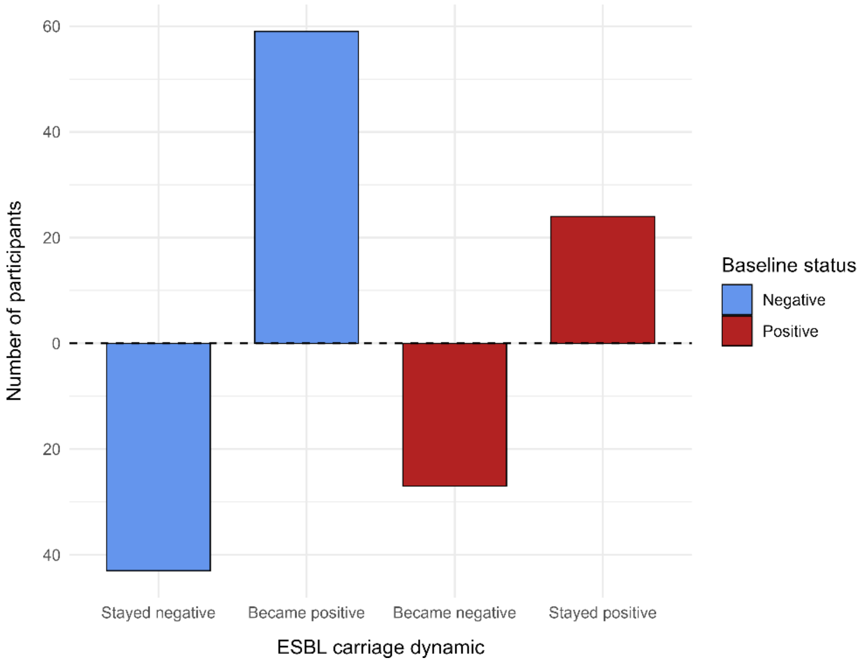

**Supplementary Figure 2. ANI hierarchical clustering dendrogram.** ANI hierarchical clustering with average linkage was used to identify strains with >99.99% ANI cutoff.

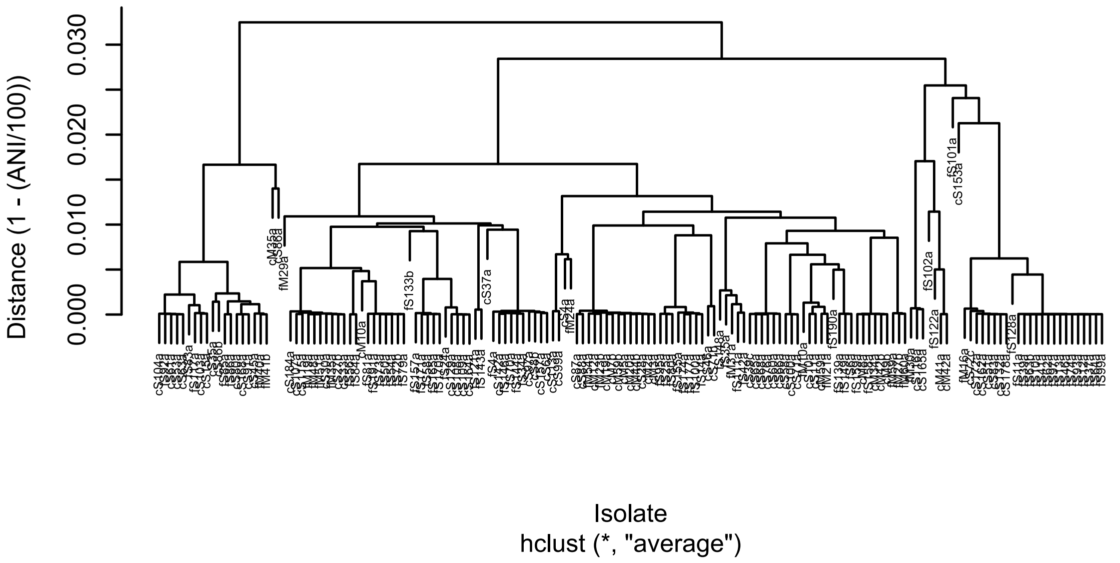

**Supplementary Figure 3. Plasmid replicon types.**

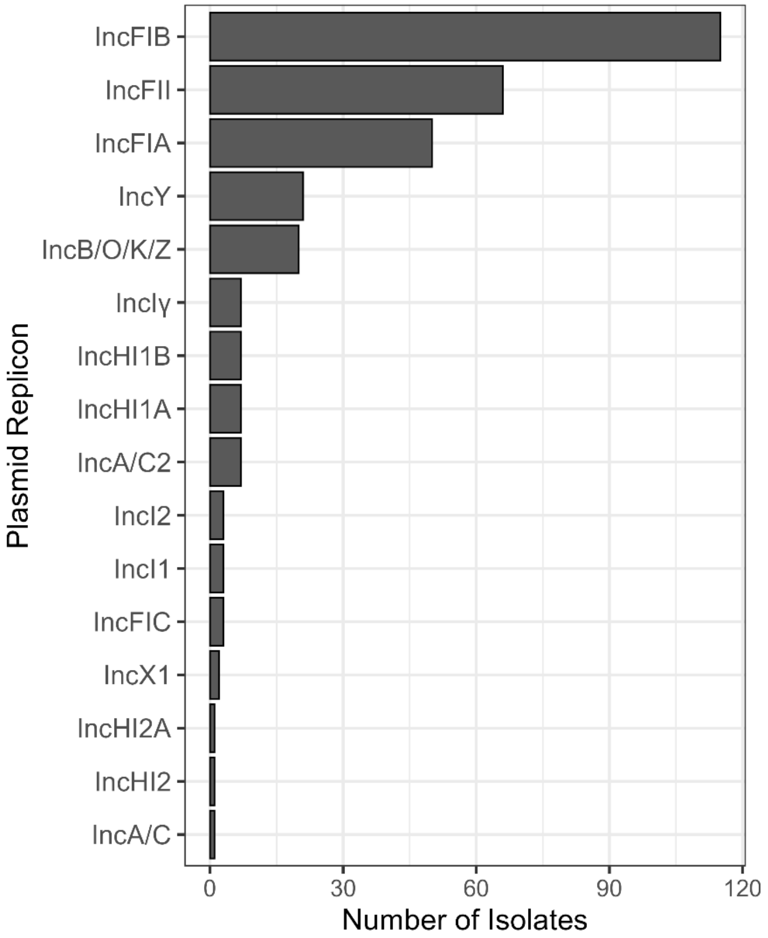

**Supplementary Figure 4. ARG profile dynamics (2023–24).**

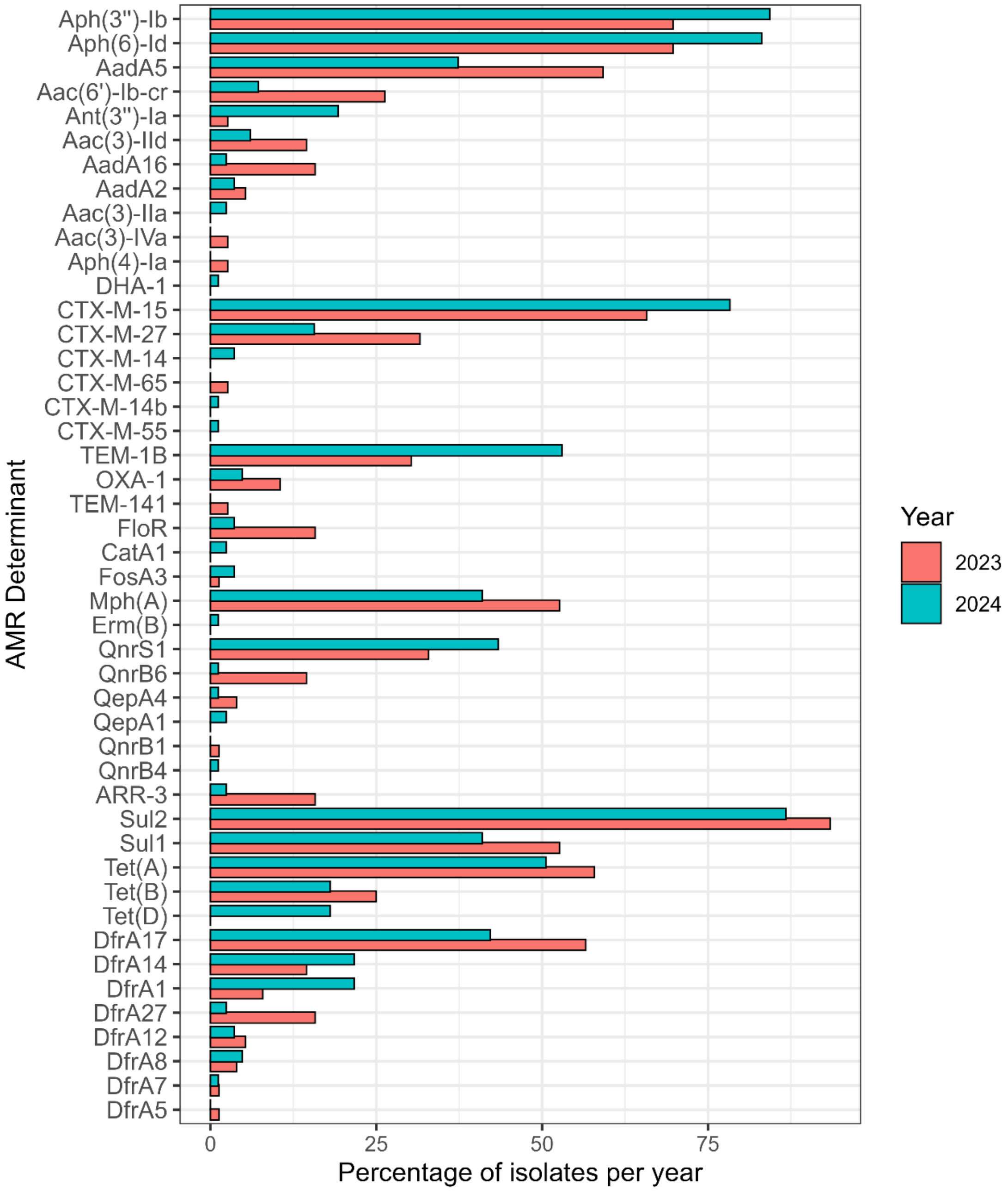

**Supplementary Figure 5. ST profile dynamics (2023–24).**

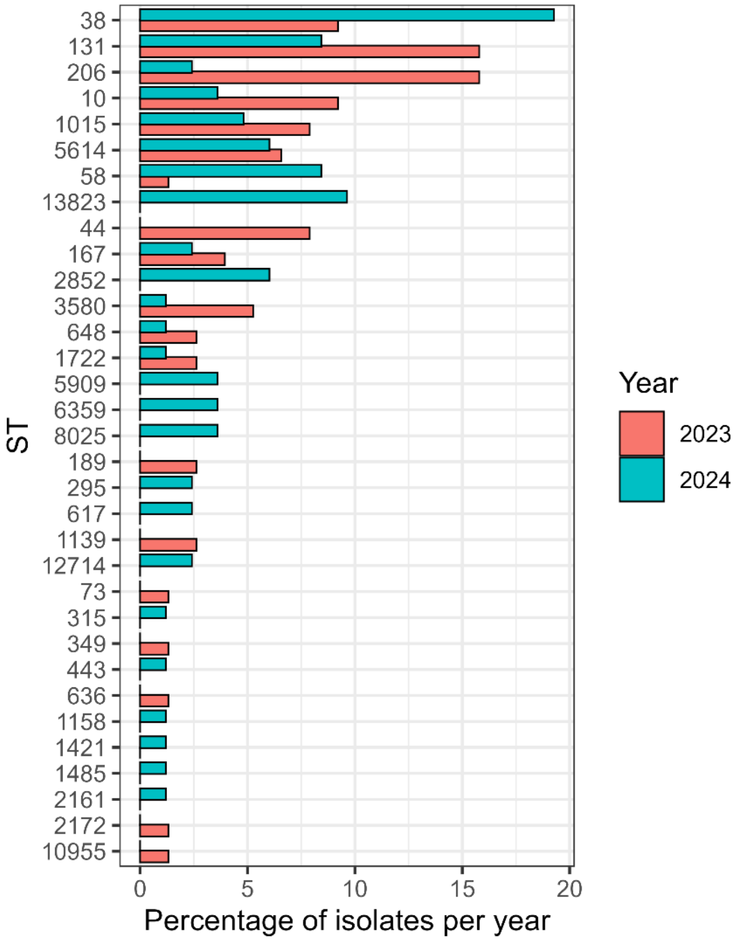

**Supplementary Figure 6. Yersiniabactin (*ybt*) biosynthetic gene cluster comparison.** Annotated medoid genomes of strains found in ≥3 households containing the *ybt* biosynthetic gene cluster are included. All links share 100% nucleotide identity. An IS*4* family transposase inserted between the *ip2* and *ybtA* genes in cM8a.

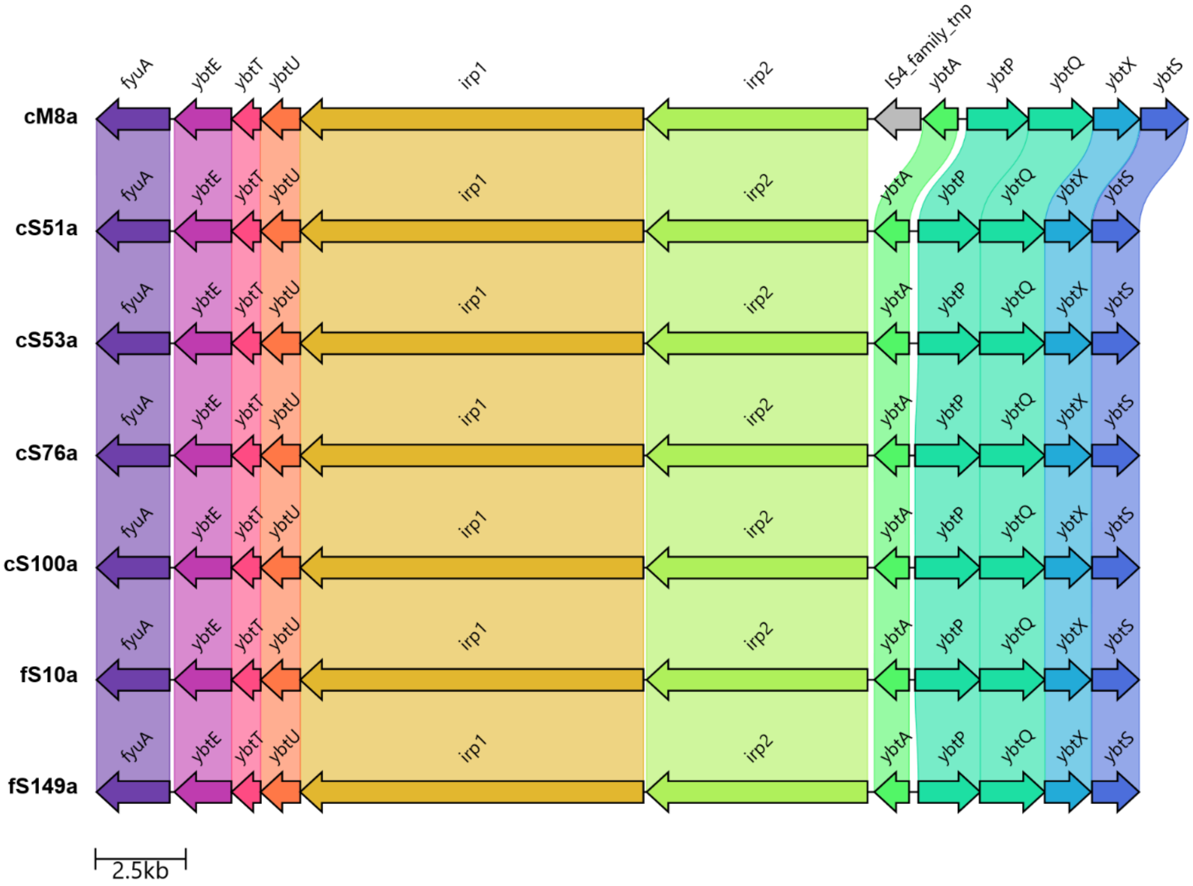

**Supplementary Figure 7. Scatter plot of pairwise SNP distances between same-strain isolates identified in ≥3 households.** Pairwise comparisons are shown between isolates sampled concurrently from different households in the same village.

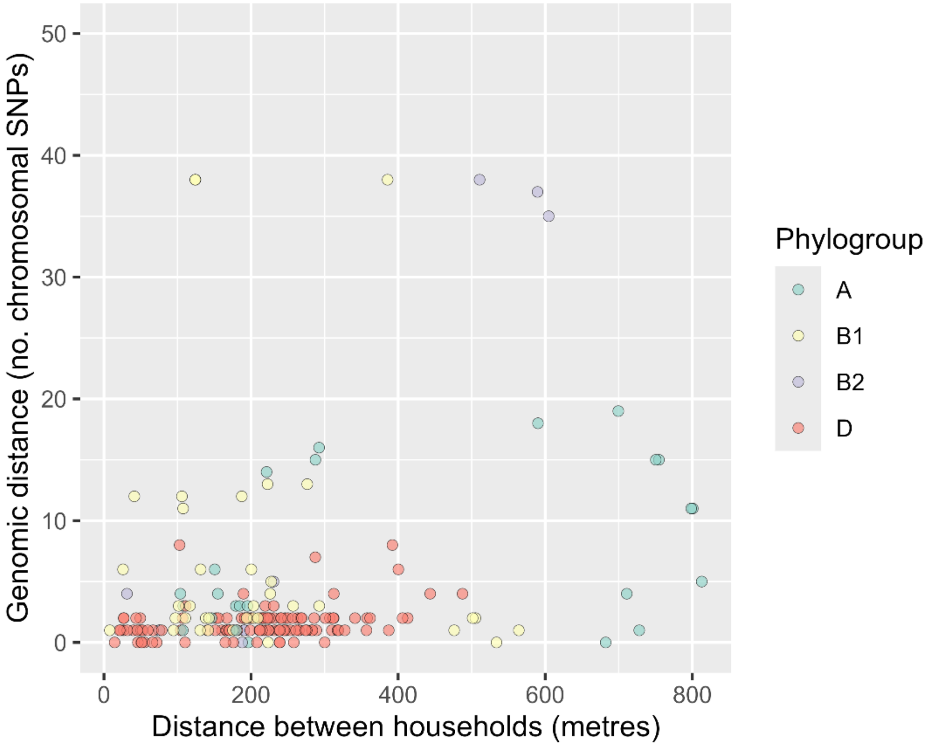
